# A myopic perspective on the future of protein diagnostics

**DOI:** 10.1101/227728

**Authors:** Ulf Landegren, Rasel A. Al-Amin, Johan Björkesten

## Abstract

Plasma proteome analyses of the future promise invaluable insights into states of health, not only by measuring proteins whose role it is to ensure blood homeostasis, but increasingly also as a window into the health of practically any tissue in the body via so-called leakage protein biomarkers. Realizing more of this vast potential will require progress along many lines. Here we discuss the main ones, such as optimal selection of target proteins, affinity reagents, immunoassay formats, samples, and applications, with a view from ongoing work in our laboratory.

---

The concept of liquid biopsy attracts interest because of the potential to improve diagnostics by revealing diseases anywhere in the body via a simple blood sampling. Cells, DNA and RNA molecules from otherwise hard-to-reach tissues can all potentially be accessed via blood samples and applied to investigate organ damage, malignancy or fetal health [1,2]. Assays for protein in plasma often target proteins that exert their activities in blood, such as coagulation factors, lipoproteins or cytokines, but liquid biopsies in the form of protein assays that target leakage markers are also well established in routine healthcare. For example, elevated plasma levels of troponin, exclusively expressed in heart muscle cells, signals insults to myocardial tissue in a heart attack [3]. Similarly, S100B is a marker of brain damage [4], possibly superseded in diagnostic value by the more recently identified serum neurofilament light protein [5].

It is likely that many more proteins than currently appreciated could provide a basis for improved diagnostics via protein-based liquid biopsy testing, and exosomes, recognized via their membrane proteins, represent a related class of targets for testing [6-9]. Affinity-based protein detection seems to offer the greatest promise for highly sensitive protein assays, but despite rapidly increasing molecular insights, progress establishing new, clinically useful markers has been surprisingly slow [10,11]. It is worthwhile taking stock of what it may take to develop and apply new protein liquid biopsy markers on a larger scale, with the purpose of assessing states of health via blood samples. Here, we consider the following questions that we see as critical for progress: What are the most relevant target proteins? What affinity reagents should be used? What assay architectures best capture this information? What samples are needed for the analyses? And how will assays be used for research and later for routine testing?

## What proteins should be targeted?

Tissue-specific proteins normally with low or undetectable levels in plasma are particularly promising as biomarker candidates, since even subtle increases may reflect damage to the tissues that express them. The challenge to identify suitable proteins that might serve as leakage markers of tissue-specific disease is greatly assisted by a number of ongoing mapping projects. The expression of genes in diverse tissues is most readily studied by sequencing their transcripts [12-14], but also the expression of proteins is being investigated via affinity- and mass spectrometry-based analyses [15,16]. Since tissues are composed of diverse cell types, including the blood cells passing through all of them, it will be necessary to look closer to find cell-specific gene expression. The recently initiated Human Cell Atlas project will provide RNA and protein expression data at the ultimate resolution of individual cells throughout the body, accounting for the diverse cell types making up all tissues [17].

Since biomarker proteins may be of particular interest for cancer diagnostics, and as most malignancies arise in cells of epithelial origin, proteins expressed by epithelial cells attract special attention. In particular, epithelial proteins destined for exocrine secretion to the lumen of the organ whose walls they line may prove particularly promising as cancer markers since these proteins are meant to leave the body, but may accumulate in blood when their normal release is obstructed [18]. Accordingly, any space-filling local process in the organ might increase blood levels of tissue specific proteins that would otherwise have been released to the lumen from low or nonexistent starting levels. Prostate specific antigen (PSA) serves to illustrate this principle, as its presence in plasma can reflect tumors or other pathological states of the prostate when its normal release to seminal fluid is prevented [19], but other proteins or exosomes may follow the same pattern.

The necessity to make educated guesses of promising proteins in the search for biomarkers will be offset by increasingly high-throughput and low-cost techniques that will allow ever broader screens for proteins of possible diagnostic value, as discussed below.

Analyses of protein biomarkers have to take into account the substantial inter-individual variation of protein levels, complicating efforts to define levels that are diagnostic for disease. This problem is being addressed by explicitly identifying factors that may account for this variation, such as genetics, age, gender, diet, etc [20,21]. In another, complementary approach, individuals can be used as their own controls for biomarker discovery, by repeatedly sampling the same individuals and observing how levels change with the emergence of ill health. This latter strategy calls for new forms of biobanks, as discussed below.

## What affinity reagents to use?

The search for leakage protein markers places stringent demands on the reagents used for their detection. For example, affinity reagents that prove suitable to detect native proteins in plasma may not recognize denatured proteins in immunohistochemistry, and vice versa [22,23]. The question whether to use polyclonal antibodies or clonal reagents such as monoclonal antibodies, recombinant affinity reagents, DNA aptamers, or maybe also drug-like, low molecular weight compounds, so far is weighted in favor of polyclonals mainly for reasons of cost. This could change with still more efficient techniques to isolate and validate clonal affinity reagents in vitro, which would simplify standardization of assays, and permit sharing of reagents that can be replenished indefinitely [23,24]. Recombinant reagents may have to be expressed in homodimeric form to achieve the avidity of natural antibodies. Whatever the source of binding reagents, there is a premium for those that bind their correct targets with high affinity.

The requirement for highly specific target detection is naturally far more demanding in assays that target tissue-specific proteins, often present at parts per billion or even less in plasma. Even if robust levels can be demonstrated for some tissue-specific proteins in plasma from patients with manifest disease, being able to detect much lower levels may translate to earlier detection of that disease. Antibodies and other affinity reagents typically exhibit demonstrable affinity for a wide range of proteins besides their intended targets, and if some of these bystander proteins are more abundant than the leakage protein of interest, as is likely to be the case, then correct signals may easily be swamped by background.

The affinity of antibodies for their targets and any cross-reactive molecules is an expression of the on- and off-rate for the antibody-target interaction [25]. The faster that antibodies are able to bind proteins, and the longer they stay bound, the higher the affinity. Some assays achieve high specificity by focusing on antibody-antigen interactions that form very rapidly. For example, in lateral flow devices each protein may have to be bound in seconds as it passes an immobilized capture antibody, thereby discriminating against nonspecific interaction with a slower on-rate [26,27]. Conversely, many assays require antibodies to remain bound to their target molecules for an extended time, again minimizing background from low-affinity interactions. This is true for typical sandwich immunoassays with long, sequential incubations and extensive washes. The emphasis on long off-rates is a crucial factor in SomaLogic Inc’s protein assays, where only those so-called slow off-rate modified aptamers used in their assays that remain bound to proteins in the sample for a considerable time can give rise to detectable signals [28]. In this manner the assays ignore weaker interactions that could have resulted in nonspecific background, thereby increasing assay specificity.

## What assay architectures provide the required performance?

Protein immunoassays can be configured so that samples are immobilized and interrogated with labeled antibodies (Fig. 1A). Alternatively, immobilized antibodies can be used to capture target molecules after labeling all molecules in the sample (Fig. 1B). Sandwich assays, by contrast, avoid the need to immobilize and/or label samples to be interrogated, since in these assays immobilized antibodies capture proteins from solution-phase samples, followed by recognition by labeled antibodies (Fig. 1C). More importantly, because in sandwich assays two antibodies must recognize each target molecule these assays offer greater specificity of detection than those where single affinity reactions suffice for detection. Only irrelevant proteins for which both antibodies exhibit cross reactivity can give rise to background signals. This greatly reduces the risk of nonspecificity and thus enables higher detection sensitivity [11].

**Fig. 1.**
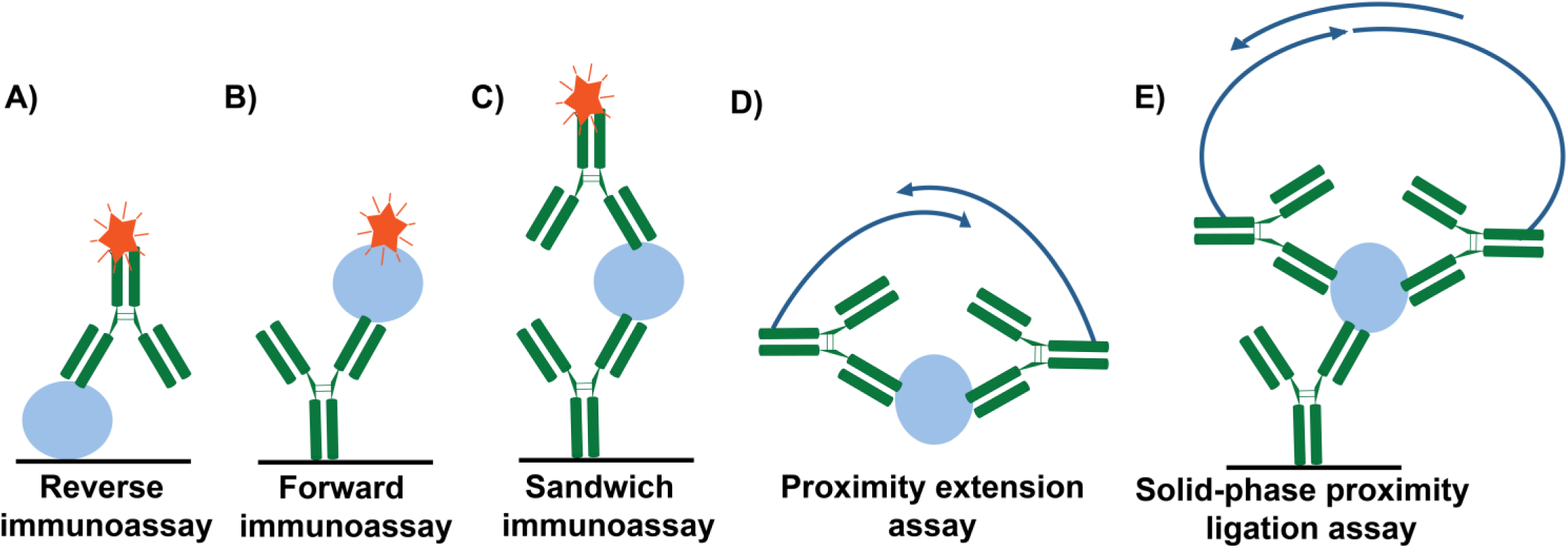
Commonly used immunoassay architectures. **A.** In reverse immunoassay proteins in immobilized samples are recognized by labeled antibodies. **B.** In forward immunoassay immobilized antibodies capture target molecules from labeled samples. **C.** In sandwich immunoassays target proteins in a sample are capture by immobilized antibodies and detected by labeled antibodies added in solution phase. **D.** In proximity extension assays target proteins in small aliquots of samples are bound by pairs of oligonucleotide-conjugated antibodies in solution phase. After an incubation the reactions are diluted to reduce chance proximity between reagents, and a DNA polymerase extends one or both oligonucleotides, templated by the other, to give rise to a reporter strand that can be quantified by realtime PCR. **E.** In solid phase proximity ligation assays an immobilized antibody captures the target protein from a sample, followed by addition of pairs of oligonucleotide-conjugated antibodies. After washes, oligonucleotides in proximity are joined by ligation to give rise to amplifiable reporter DNA strands, quantifiable by realtime PCR.

Besides cross-reactive detection of irrelevant proteins in a sample, nonspecific signals also commonly depend on background in the form of some level of e.g. fluorescence or optical absorption by solutions and vessels used, depending on the readout. This source of background can be avoided entirely by designing assays where only the specific detection reagents are capable of giving rise to detectable signals. The companies Singulex Inc. and Quanterix Corp. achieve high sensitivity by developing sandwich assays where detection reactions are divided into volume elements so small that only specifically labeled antibodies give rise to signals that exceed detection thresholds [29,30]. Similarly, immune PCR and immune RCA reactions exploit DNA conjugated detection reagents that give rise to detectable amplification products, which cannot arise in their absence, thereby avoiding any contribution to detection signals by factors other than specifically or nonspecifically bound detection probes [31-33]. We are thus left with one further source of nonspecific signals, namely failure to remove labeled affinity reagents due to sticking to the reaction vessel or similar. A combination of the various sources of background, together with difficulties of recognizing more than a fraction of all target molecules in a sample, have so far conspired to keep detection of every individual protein molecule in a sample firmly out of reach for contemporary protein assays.

Similar in character to sandwich immune assays, the proximity ligation assay (PLA) developed in our laboratory, and the related proximity extension assay (PEA), commercialized by Olink Proteomics, also achieve high specificity by using pairs of antibodies or other affinity reagents for each targeted protein (Fig. 1D). For proximity assays these affinity reagents are modified by being conjugated to DNA strands. Upon pairwise binding to target molecules, DNA sequence elements that serve to identify the affinity reagents are combined into a single DNA strand through ligation or polymerization reactions in PLA and PEA reactions, respectively [34-36]. These DNA strands are then quantified by real time PCR or through DNA sequencing. The assays do not employ solid supports; instead, signals arising from chance proximity between pairs of affinity reagents are minimized by diluting the reactions for single microliter samples after incubation with detection reagents, with the addition of ligases or polymerases. This provides for a convenient homogenous assay protocol and sensitive protein detection. The assays are combined in multiplex detection reactions for around 100 target proteins without the loss of specificity normally seen in multiplex sandwich detection reactions, since only DNA strands that form in reactions between the relevant pairs of antibodies are recorded as true signals, ignoring any products from noncognate reagent pairs.

In a variant of these two proximity techniques, even higher detection sensitivity may be achieved by first capturing target molecules from a larger sample volume via immobilized antibodies, before the addition of the pair of oligonucleotide-conjugated antibodies (Fig. 1E)[37,38]. This is a more laborious procedure compared to the homogenous forms of proximity assays, and more similar to a regular sandwich ELISA. Advantages include the possibility to search for weakly expressed proteins in larger sample volumes; excess reagents may be removed by washes to reduce background; and each protein has to be recognized by a total of three antibodies, further reducing the risks of detecting off-targets through cross-reactive binding. Moreover, since only pairs of detection reagents binding in proximity, but not individual reagents, can give rise to detectable signals, risks of background from nonspecifically bound reagents are further reduced.

## What samples should be collected for analysis of protein markers?

The search for protein biomarkers that permit detection of disease at early, hopefully still curable stages, will depend not only on suitably high-performance reagents and assays, but appropriate prospective samples, collected before onset of disease and complemented by high-quality health records and results from other molecular analyses are also critical factors. As already discussed, collections of consecutive samples from the same individuals will provide a valuable opportunity to measure trends for the levels of the biomarker candidates in plasma over time, whereby individuals serve as their own controls. The availability of longitudinal samples for very large groups of patients will also increase the probability that sufficient numbers of samples will have been collected at a time when detection of biomarkers leading to a correct diagnosis could still allow abrogation of disease, although the initial discovery of such markers may be easier by focusing on more advanced cases of disease.

It is well known that the precise conditions for collecting and storing blood or plasma samples can seriously impact the results of protein assays [39,40], and simple, standardized procedures that can be applied on a large scale are an important aim in biobanking.

In this regard, we have recently demonstrated that a 1.2 mm diameter disk of paper, punched from a dried blood spot, can be interrogated by multiplex PEA with proteins equally detectable and results as reproducible as those from fresh blood samples, but with some deviations of recorded levels comparing wet and dry samples (Fig. 2). We furthermore found that the levels of most proteins remained stable upon storage for tens of years [41]. For those proteins whose levels showed a tendency to slowly decrease over time, storage at −24°C better preserved levels than did +4°C This means that very large biobanks can be built at little cost by simply drying drops of blood on paper and preserving these cold. The samples can be retrieved from standard blood samples, rather than discarding these after the prescribed analyses have been completed in healthcare. It is also possible to have individuals themselves prick their fingers, and collect the blood on papers that may be sent by regular mail for storage and analysis. The simplicity and low cost of dried blood biobanks can thus help ensure that samples will be available for biomarker discovery at critical phases of the course of disease.

**Fig. 2.**
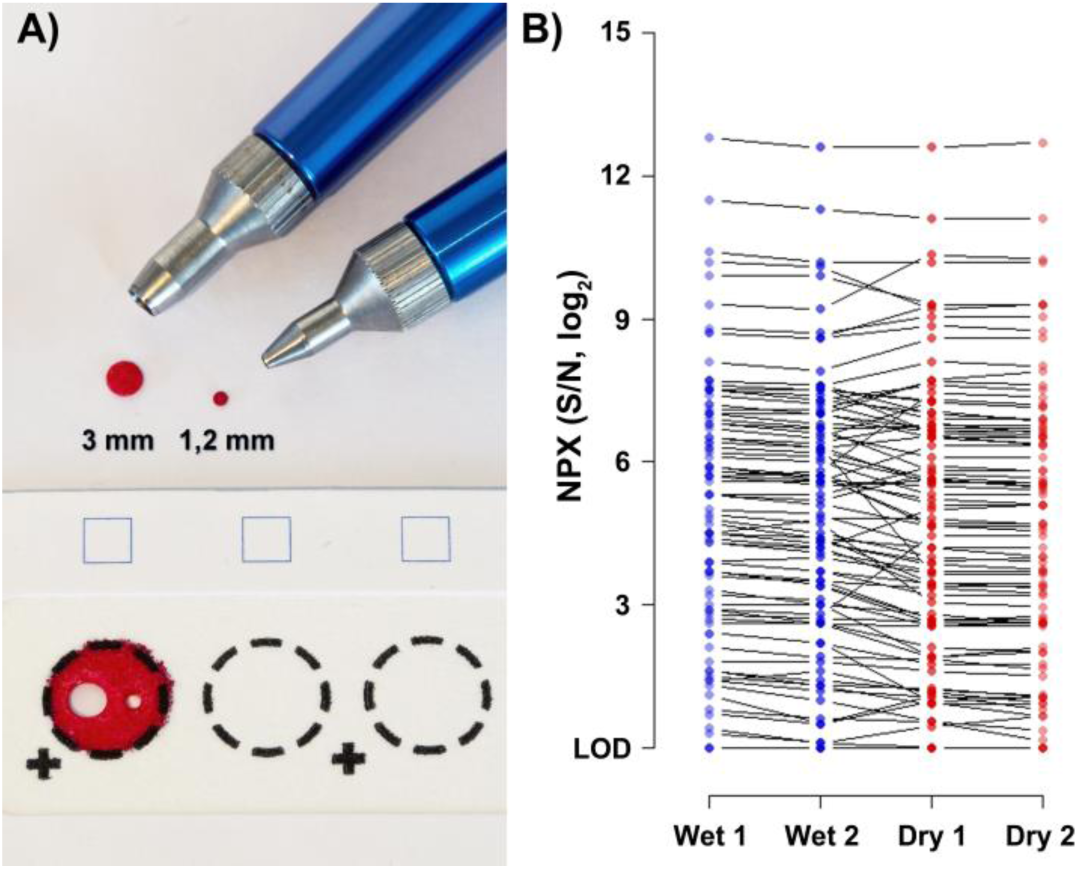
Dried blood spot sample collection. **A**. Only very small amount of dried blood spot samples are required to interrogate by multiplex PEA. The smaller 1.2 mm diameter dot suffices for sensitive analysis of sets of 92 proteins. **B**. Results from a comparison of analyses of sets of 92 protein markers in 1 μl blood, or in 1.2 mm diameter dots cut from dried blood spots on paper. Duplicate wet or dried blood samples (blue and red, respectively) were compared with each other (results replotted from reference 36).

## In what contexts will protein assay be used?

The biomarker candidates, affinity reagents, molecular technologies and samples that are collected all need to be implemented in suitable assay formats for analysis of protein leakage markers. The appetite for inexpensive, precise parallel protein analyses is likely to continue to grow, and the quality requirements will become more stringent compared to the current sometimes relatively lax standards for home-brew assays used in research labs. The volume of protein assays in research will expand greatly with the screening of increasing numbers of proteins in rapidly growing sample collections. As knowledge builds, applications of protein assays are sure to grow in importance also for drug development, in clinical diagnostics, and for wellness monitoring where the aim is to measure factors that can help maintaining states of health [42].

Costs for genotyping analyses at the level of DNA in research have gone from $1 per single nucleotide polymorphism (SNP) to around 100,000 SNPs to the $ since the beginning of the millennium. High-throughput protein assays in research are currently priced somewhere around $1 per protein and sample, but a similar price erosion as for DNA analyses is possible for protein measurements as well, commensurate with a vast increase in the numbers of analyses to be undertaken in the search for new biomarkers. To achieve this cost reduction, critical elements of the assays have to be scrutinized, such as affinity probes, enzymes and other reagents, disposables and the technique used for readout of assay results. There is also a need to continue improving study design and evaluation of the significance of the results.

In clinical diagnostics, moderate size panels of plasma protein markers may allow finer distinctions between different states of disease than what is possible with individual markers. As discussed there is also great hope that analyses of leakage markers will allow disease to be discovered at presymptomatic stages by regular screens of broad populations at risk due to factors such as age, gender or life style via dedicated marker panels. Quality requirements for diagnostic tests are substantially more stringent than those that apply in research, but requirements for turn-around time in general can still be modest.

As a final point, emerging tests for proteins that can reflect the integrity of tissues will increasingly also be formatted for rapid applications at the point of care, sometimes with modest levels of multiplexing. Yet another area of significant growth outside healthcare is that of wellness testing, where blood from finger pricks may be sent for updates about the day to day health of organs, perhaps serving to guide the choice of diet and exercise, etc, and only exceptionally prompting contacts with standard healthcare. All in all, the volumes of protein assays are likely to increase dramatically, and so will in all likelihood the value of protein assays as important means to maintain health and to identify disease at the earliest possible time.

## Disclosure

UL is founder, shareholder, and board member of Olink Proteomics, commercializing proximity extension assays.

## Highlights

1. Protein leakage markers can allow liquid biopsies that reveal disease processes
2. Leakage markers promise diagnosis at early timepoints, but progress has been slow
3. Tissue-specific proteins are of special interest as potential leakage markers
4. High-quality affinity reagents remain a limiting factor for new assays
5. Target recognition by two or more antibodies improves specificity
6. Extensive biobanks of dried blood spot could serve to validate markers
7. Greatly increased protein assay throughput can be foreseen in research

## Acknowledgements

Work in the authors’ laboratory is funded by the European Community’s 7th Framework Program (FP7/2007-2013) under grant agreement n° 313010 (BBMRI-LPC), the Swedish Research Council, the IngaBritt and Arne Lundberg’s Research Foundation, Torsten Söderberg’s Foundation, and the Swedish Foundation for Strategic Research.

